# Hydra Bcl-2 and TMBIMP family proteins display anti-apoptotic functions

**DOI:** 10.1101/359174

**Authors:** Mina Motamedi, Laura Lindenthal, Anita Wagner, Margherita Kemper, Jasmin Moneer, Mona Steichele, Alexander Klimovich, Jörg Wittlieb, Marcell Jenewein, Angelika Böttger

## Abstract

**Background:** Mechanisms of programmed cell death differ considerably between animals, plants and fungi. In animals they depend on caspases and Bcl-2 family proteins and this kind of cell death is called apoptosis. Most gene families encoding proteins involved in apoptosis are found in multicellular animals already in the eldest phyla but their functional conservation is still being studied. Much older protein families have cytoprotective functions across all kingdoms of life. This includes the TMBIMP-family, the presence and function of which in early metazoans has not been investigated yet.

**Methods:** We quantified apoptosis in transgenic *Hydra* overexpressing HyBcl-2-like 4. Moreover, we investigated putative TMBIMP-family members in *Hydra* by sequence comparison. By overexpression of TMBIMP-family members in *Hydra* and human HEK cells we analysed their subcellular localisation and in one case their capacity to protect cells from camptothecin induced apoptosis.

**Results:** HyBcl-2-like 4, as previously shown in a heterologous system, was localised to mitochondria and able to protect *Hydra* epithelial cells from apoptosis. The TMBIMP-family in *Hydra* includes HyBax-Inhibitor-1, HyLifeguard-1a and -1b and HyLifeguard 4 proteins. HyBax-inhibitor-1 protein was found localised to ER-membranes, HyLifeguard-family members were found at the plasma membrane and in Golgi-vesicles. Moreover, HyBax-inhibitor-1 protected human cells from apoptosis.

**Conclusion:** This work provides the first functional study to support an anti-apoptotic function of Bcl-2 like proteins in pre-bilaterians within a physiological context. Furthermore it illustrates that genes that were “inherited” from non-animal ancestors, like the TMBIMP-family, were recruited to carry out cell protective anti-apoptotic functions already in early metazoans.

## Introduction

Programmed cell death has been observed in almost all organisms including protozoans, fungi, plants and animals (Falcone and Mazzoni, 2016; Kaczanowski et al., 2011; Reece et al., 2011; Van Durme and Nowack, 2016). It is crucial for removal of damaged cells and functions as an important regulator of development.

The molecular mechanisms governing induction, regulation and execution of programmed cell death differ between the different kingdoms. In animals, they involve the activation of cystein proteases (caspases) through aspartate specific substrate cleavage. In vertebrates, including humans, caspase activation proceeds either via extrinsic receptor-based stimuli or through intrinsic pathways, which employ members of the Bcl-2 protein family. Programmed cell death that depends on caspases and Bcl-2 proteins is commonly referred to as apoptosis and is only found in animals. In mammals, most Bcl-2 proteins are localised at the mitochondrial outer membrane and only a few are found associated with ER-membranes. Upon apoptosis-inducing signals, the outer mitochondrial membrane gets permeabilised by factors that inhibit the anti-apoptotic Bcl-2 family proteins. At the same time several pro-apoptotic factors, including cytochrome C, are released from the intra-membrane space. The cytosolic cytochrome C then forms an apoptosome with APAF-1 and initiator caspases, thereby promoting their self-activation and subsequent activation of effector caspases, which cleave specific substrates to induce all the morphological and molecular changes that lead to cell death.

The Bcl-2 gene family is conserved from early invertebrates, including *Caenorhabditis* and *Drosophila*, to humans. However, their functions for intrinsic apoptosis induction differ between invertebrates and vertebrates. In *Caenorhabditis* the BH-3-only protein Egl-1 induces apoptosis via inhibition of ced-9 (the only Bcl-2 homolog found in *Caenorhabditis*), release of ced-4 (APF-1) and activation of ced-3 (caspase 3) (Conradt and Horvitz, 1998). The role of the two *Drosophila* Bcl-2 family members, *debcl* and *buffy*, in apoptosis is unclear. Both genes are not essential for the development of flies.

To better understand the evolution of apoptosis we have previously analysed caspases and Bcl-2 family members in the pre-bilaterian cnidarian model organism *Hydra* (Lasi et al., 2010a; Lasi et al., 2010b). In this context we had originally identified seven Bcl-2-like and two Bak-like genes. Additional members of the *Hydra* Bcl-2 family were found through cell type specific transcriptome analyses (Wenger et al., 2016). In accordance with these published data a recent search in available sequencing data from *Hydra* has now revealed 9 genes encoding Bcl-2-like proteins and two encoding Bak-like proteins. This gene family was also found recently to be present in corals of the genus *Acropora* ((Moya et al., 2016)). The *Hydra* proteins that we had previously studied were associated with membranes, five with mitochondria- and two with ER-membranes. When expressed in HEK (human embryonic kidney) cells three family members induced apoptosis. The other Bcl-2 like proteins protected HEK-cells from camptothecin induced apoptosis. Moreover, the *Hydra* genome encodes several BH-3 only proteins. Of those, a 96 amino acid protein, which we named HyBH3-only 2, induced apoptosis in human cells (Lasi et al., 2010b). Collectively, these data indicate that the genes governing intrinsic apoptosis-inducing pathways involving mitochondria, as we find them in mammals, have appeared before the evolution of bilaterian animals.

Apoptosis in Hydra is found at the extremities of the animals, where cells are sloughed off and replaced by new ones. Moreover, it is used to regulate cell numbers under conditions of variant nutrient supply ((Bosch and David, 1984) and reviewed in (Böttger and Alexandrova, 2007; Reiter et al., 2012)). The role of Bcl-2 proteins in these developmental processes in *Hydra* is not understood in much detail yet. In this work we present the first functional study of a Bcl-2 protein in cnidarians. We investigated a transgenic *Hydra* line overexpressing HyBcl-2-like 4 protein in mitochondria of ectodermal cells. We found that the elevated levels of HyBcl-2-like 4 in this *Hydra* line protected *Hydra* epithelial cells from Wortmannin and starvation induced apoptosis.

In contrast to the Bcl-2 protein family, which has only been detected in animals, the family of “**t**ransmembrane **b**ax-inhibitor **m**otif **p**roteins” or TMBIMPs, is conserved in animals, protists, plants and fungi (Hu et al., 2009). Members of this membrane protein family are cytoprotective and implicated in the regulation of programmed cell death. The name giving and founding member of the family is Bax-Inhibitor (BI-1 or TMBIMP6), (Rojas-Rivera and Hetz, 2015). BI-1 plays a role in protection from ER-stress, including such caused bydisturbances in Ca^2+^-homeostasis and unfolded protein response (reviewed in (Ishikawa et al., 2011)). The neuronal membrane protein TMBIMP 2 obtained the name “life guard” (Lfg-2) due to its ability to inhibit FasL induced apoptosis (Somia et al., 1999). This name was then applied to other TMBIMP family members including Lfg-1, -3, -4 and -5. Lfg-4 is a Golgi-protein, which protects cells against both intrinsic and extrinsic apoptosis activation (de Mattia et al., 2009; Gubser et al., 2007). Lfg-3, when deleted, increases susceptibility of mice to medial cystic degeneration (Zhao et al., 2006). Whether Lfg-1 and Lfg-5 are also involved in the regulation of apoptosis is not known to date (summarised in (Hu et al., 2009). A recent study by Lisak et al (Lisak et al., 2015) investigated the subcellular distribution of all TMBIM family members in human cells. They found that TMBIM 1, 2 and 3 (corresponding to Lfg-3, - 2 and -1) were localised in Golgi vesicles whereas TMBIM 4, 5 and 6 (corresponding to Lfg-4, Lfg-5 and BI-1) were localised at the ER. Lfg-5 (TMBIM5) was special, because it has an N-terminal signal sequence that can target it to mitochondria. An N-terminally tagged version of this protein localised to the ER.

Here we have studied the TMBIMP-protein family in *Hydra*. We have cloned four genes encoding HyBI-1 and three Lfg-like proteins. According to their phylogenetic relationships we named them HyLfg-1a and -1b and HyLfg-4. We found HyBI-1 localised at the ER whereas the three HyLfg-like proteins were associated with the Golgi apparatus. HyBI-1 exhibited an apoptosis protective function in human cells.

## Material and methods

### Hydra culture

*Hydra vulgaris* strain Basel was cultured at a constant temperature of 18 °C in hydra medium (0.1 mM KCl, 1 mM NaCl, 0.1 mM MgSO_4_, 1 mM Tris and 1 mM CaCl_2_). The animals were fed regularly with freshly hatched *Artemia nauplii* from Sanders Brine Shrimp Company. All animals were starved at least 24 hour prior to be used in experiments.

### Gene Cloning

A search of whole genome and EST (Expressed sequence tag) sequences from *Hydra* (http://hydrazome.metazome.net/cgi-bin/gbrowse/hydra/) has revealed three sequences with homology to human TMBIMP-family members. Through Blast analysis, *HyLfg-4* (XM_002162893.1) and two Lfg-1 homologues were found in *Hydra*. They are named *HyLfg-1a* (Hma2.214458) and *HyLfg-1b* (Hma2.205245) in this study. Based on gene models for each homolog of Lifeguard in *Hydra*, pairs of primers were designed. These primers were used for PCR amplification from *Hydra* cDNA and cloning of the respective sequences into the HoTG expression vector using the SmaI restriction site (Böttger et al., 2002). Another member of the TMBIMP family, *HyBI-1*, had previously been identified in the *Hydra* genome (Hma1.130444) (Lasi et al., 2010).

### Transfection of *Hydra* cells and transgenic animals

Gold particles (1.0 μm, BioRad) were coated with plasmid DNA according to instructions of the manufacturer. They were introduced into *Hydra* cells with the Helios gene gun system (BioRad) as previously described (Böttger et al., 2002). For transgenic animals HoTG plasmids with the *Hydra* actin promoter encoding GFP-HyBcl-2-like 4 and mitochondrially targeted GFP were injected into fertilised eggs as previously described (Wittlieb et al., 2006). After development of the embryos, transgenic animals were detected and it was continuously selected for animals with strongly fluorescing mitochondria. In this way the two lines we investigated in study were obtained.

### Confocal Laser Scanning Microscopy

Leica SP5-2 confocal laser-scanning microscope was used for Light optical serial sections. Leica SP5-2 equipped with an oil immersion Plan-Apochromat 100/1.4 NA objective lens. EGFP, FITC, Alexa488 were visualized with an argon laser at excitation wavelength of 488 nm and emission filter at 520 - 540 nm for eGFP, FITC and Alexa488. The helium-neon laser with excitation wavelength of 633 nm and emission filter of 660–760 nm was used for TO-PRO-3. Cy3, RFP and BODIPY-TR were visualized using a Krypton laser excited at a wavelength of 561nm and emission filter at 570-580nm. Image resolution was 512 × 512 pixel with a pixel size ranging from 195 to 49 nm depending on the selected zoom factor. The axial distance between optical sections was 300 nm. To obtain an improved signal-to-noise ratio, each section image was averaged from three successive scans. The 8-bit grey scale single channel images were overlaid to an RGB image assigning a false colour to each channel and then assembled into tables using Adobe Photoshop 8.0.

### Confocal microscopy on living animals

In a live-scan, the animals were initially relaxed with 2% urethane for 2-3min and placed on a slide with a wax-foot coverslip. For additional staining with the Golgi marker, 1μl of 500μM BODIPY-TR in hydra medium solution was injected into the hydra gastric cavity. Thereafter, polyps were incubated in 5μM BODIPY-TR ceramide solution for 30min. Immediately afterwards the optical sections were made.

### Phylogenetic tree

The phylogenetic tree was created based on Maximum likelihood tree (ML). Muscle alignments with 10,000 bootstrap replications were used. Nucleotide accession number list: *Hydra vulgaris* lfg4: XM_002162893.3; *Hydra vulgaris* lfg1b: XM_012709043.1; *Hydra vulgaris* lfg1a: XM_012704850.1; *Hydra vulgaris* BI1: XM_002159829.3; *Nematostella vectensis* lfg4: XM_001640172.1; *Nematostella vectensis* Lfg1b: XM_001641388.1; *Nematostella vectensis* Lfg1a: XM_001629587.1; *Nematostella vectensis* BI1: XM_001637805.1; *Xenopus tropicalis* lfg4: NM_001004879.1; *Xenopus tropicalis* Lfg3: NM_001030497.1; *Xenopus tropicalis* Lfg2: NM_001078889.1; *Xenopus tropicalis* Lfg1: NM_001016038.2; Pan troglodytes Lfg5: XM_016957765.1; *Arabidopsis thaliana* Lfg4a: NM_100189.2; *Arabidopsis thaliana* Lfg4b: AY735633.2; *Arabidopsis thaliana* Lfg4c: NM_116196.2; *Arabidopsis thaliana* Lfg4d: NM_117558.2; *Arabidopsis thaliana* Lfg4e: NM_117636.2; *Arabidopsis thaliana* BI1a: NM_117865.1; *Arabidopsis thaliana* BI1b: NM_124084.1; *Arabidopsis thaliana* BI1c: NM_117636.2; *Bos taurus* Lfg5: NM_001077068.1; *Homo sapiens* Lfg5: NM_001317905.1; *Homo sapiens* Lfg4: NM_016056.3; *Homo sapiens* Lfg3: NM_001321429.1; *Homo sapiens* Lfg2: XM_017019040.1; *Homo sapiens* Lfg1: NM_001009184.1; *Homo sapiens* BI-1: AY736129.1; *Ceratopteris richardii* lfg4: CV734669; *Danio rerio* lfg4: BC057432.1; *Danio rerio* lfg3: BC083414.1; *Danio rerio* lfg2: NM_001013518.1; *Danio rerio* lfg1: BC053253.1; *Danio rerio* BI-1: XM_009305712.2; *Chlamydomonas reinhardtii* lfg: DS496110.1; *Physcomitrella patens* Lfga: XM_001756985.1; *Physcomitrella patens* Lfgb: XM_001768570.1; *Physcomitrella patens* Lfgc: XM_001757644.1; *Mus musculus* Lfg5: NM_029141.4; *Mus musculus* Lfg4: NM_026617.3; *Mus musculus* Lfg3: NM_027154.5; *Mus musculus* Lfg2: NM_028224.4; *Mus musculus* Lfg1: XM_006521251.3; *Mus musculus* BI1: NM_026669.4; *Cyanidioschyzon merolae* lfg4: AP006486 chromosome 4: 54288-55103; *Drosophila melanogaster* BI1: NM_139948.4; *Zea mays* Lfga: BT088363.1; *Zea mays* Lfgb: NM_001156405.2; *Zea mays* Lfgc: NM_001137112.1; *Zea mays* Lfgd: NM_001321409.1; *Canis lupus* Lfg5: XM_847212.4; *Xenopus laevis* BI1: NM_001087329.1; *Picea glauca* lfg4a: EX444921.1; *Picea glauca* lfg4b: EX425899.1; *Picea glauca* lfg4c: EX375038.1; *Gallus gallus* lfg4: XP_001235093.1; *Gallus gallus* lfg3: XM_422067.4; *Gallus gallus* lfg2: XM_424507.3; *Caenorhabditis elegans* lfg4: NM_077142.5; *Caenorhabditis elegans* lfg1a: NM_068949.3; *Caenorhabditis elegans* lfg1b: NM_073099.5; *Saccharomyces cerevisiae* lfg: NM_001183143.1 *Ovis aries* Lfg5: XM_012176315.2. Genious.8 phylogeny software was used for tree drawing.

### Apoptosis assay in *Hydra*

Animals were kept aside from the main culture for 2 days, and then incubated with four different concentrations of Wortmannin [1167 nM, 116.7 nM, 11.67 nM, 0 nM – each in 1% DMSO] for 4.6 hours at 18 °C. Afterwards, the animals were macerated with 270 µl maceration solution (glycerol, glacial acetic acid, H2O – 1:1:13 – v/v/v) for 1 hour at room temperature, and subsequently incubated for 10 minutes at room temperature in an equal volume of 8% PFA (paraformaldehyde) in PBS (phosphate buffered saline). The cell suspension was transferred with approximately 50µL 0.1% Tween onto a defined area on a microscope slide, and dried for one hour under the hood. The dried slide was washed twice with 1x PBS for 10 minutes and incubated with 50µL DAPI [1µg/µL] for four minutes. After two more washes with PBS for 10 minutes each, they were mounted with Vectashield and sealed with transparent nail polish. Under the fluorescence microscope, 1000 epithelial cells were counted per time point and concentration, and the number of apoptotic cells was determined on the basis of the apoptotic morphology of the DAPI-stained nuclei (Lasi et al., 2010b). Results are from four biological replicates.

### Statistical analysis

The statistical analysis was implemented using the program SigmaPlot 11.0, a Two-Way ANOVA (Tukey test) was applied to the Wortmannin treated *Hydra* (Fig. 2A), a One-Way ANOVA (Tukey test) was applied to the starved *Hydra* (Fig. 2B). The significance level α was set to 0.05, which means that p-values ≤ 0.05 were assumed statistically significant and are indicated by letters. The same letters on top of the columns indicate no significant differences - differing letters indicate significance.

### Apoptosis assay in HEK293T cells

Apoptosis assays in HEK293T-cells were performed as described by Lasi et al (Lasi et al., 2010b). Briefly, HEK293T-cells were transfected with plasmids encoding HyBI-1 or HyLfg-4. After 24 hours apoptosis was induced with 10µM camptothecin. Transfected cells were counted according to their GFP-expression by observing them with a fluorescence microscope. The percentage of apoptotic cells expressing GFP or the respective fusion proteins was estimated on the basis of changes in morphology (including fragmentation and condensation of chromatin) in DAPI stained nuclei.

### Expression studies using quantitative RT-PCR

Total RNA was isolated from *Hydra magnipapillata* using the Quiagen RNeasy^®^ Plus Mini kit. The concentration and quality of the tRNA was determined using Agilent’s 2100 Bioanalyzer. Total RNA with a RNA Integrity Number > 8 was reverse transcribed using the iScript™ cDNA Synthesis Kit (Bio-Rad). The validation experiments were performed using SYBR^®^ Select Master Mix for CFX (Life Technologies) on a CF™ thermocycler (Bio-Rad) with white plastics for enhanced detection.

## Results

### *Hydra* Bcl-2-like 4 protects from apoptosis induced by Wortmannin or nutrient deprivation

Some members of the extended Bcl-2-like protein family in *Hydra* are able to protect human cells from camptothecin induced apoptosis in heterologous assays (Lasi et al., 2010b). The strongest protective effect was observed with the mitochondrially localised Bcl-2-like 4. To gain insight into the physiological role of this protein we investigated its effect on apoptosis in *Hydra*. Therefore, we analysed transgenic animals overexpressing a GFP-HyBcl2-like 4-fusion protein in the mitochondria of all ectodermal cells. These had been obtained by injecting the HoTG-plasmid encoding this protein under the control of the *Hydra* actin promoter into *Hydra* oocytes. For control we used animals that expressed mitochondrial GFP in ectodermal cells (Müller-Taubenberger et al., 2006; Wittlieb et al., 2006), obtained by injecting the plasmid HoTG:mitoGFP encoding the mitochondrial targeting sequence of *Hydra*AIF in frame with GFP. Fig. 1 shows a live image of the cytoplasm of a *Hydra* ectodermal epithelial cell overexpressing GFP-HyBcl-2-like 4. The GFP signal clearly localises to the outer part of mitochondria in contrast to MitoTrackerRed CMXRos, which accumulates in the whole of active mitochondria. This suggests that HyBcl2-like 4 is present in the outer mitochondrial membrane.

**Figure 1:**
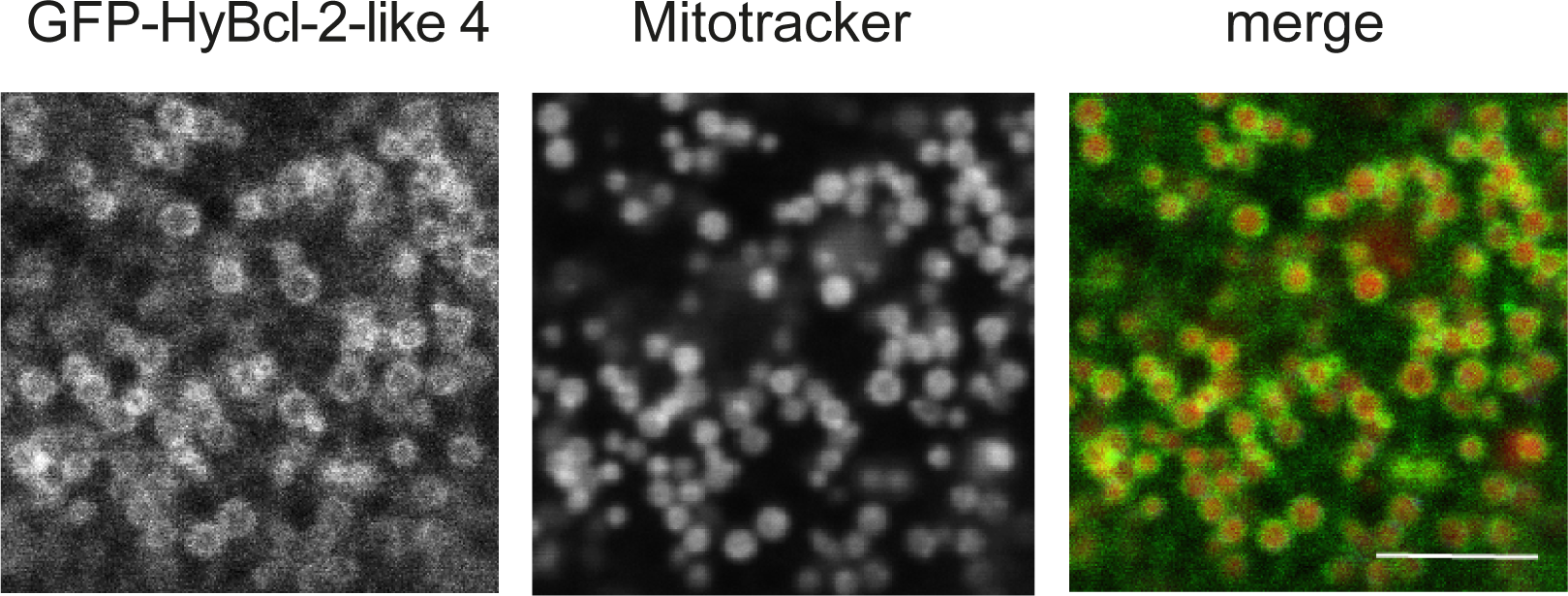
Confocal microscopic optical section from the ectoderm of the body column of transgenic *Hydra* polyps expressing GFP-HyBcl-2-like 4 in ectodermal epithelial cells. Co-staining with mitochondrial marker Mitotracker RedCMXRos (Invitrogen). Scale bars 5µm

Apoptosis was induced with the PI(3) kinase inhibitor Wortmannin. Fig. 2A shows that Wortmannin induced apoptosis in *Hydra* epithelial cells in a concentration dependent manner (apoptotic cells of the interstitial lineage were not considered). However, apoptosis was significantly reduced in HyBcl-2-like 4-GFP transgenic animals compared with mitoGFP-control animals and non-transgenic control animals.

**Figure 2:**
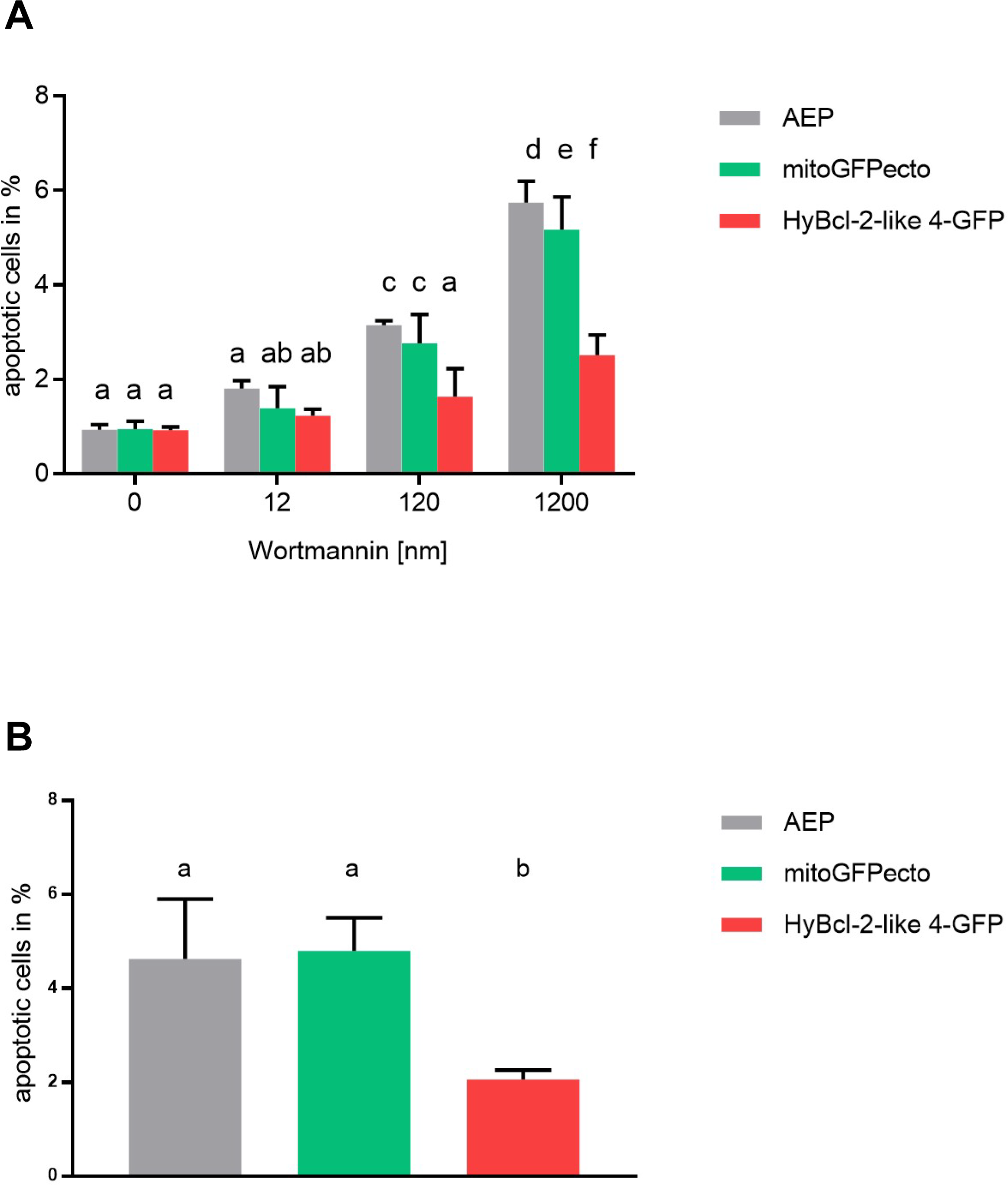
A: Diagram representing percentage of apoptotic *Hydra* cells relative to numbers of epithelial cells after treatment of polyps with PI(3)-kinase inhibitor Wortmannin. AEP founder strain (grey) and mitoGFPecto (green) animals were used for control. Standard deviation from counts of four biological replicates ; letters above columns indicate significant differences between numbers of apoptotic cells at different concentrations of Wortmannin and in different *Hydra* lines. The same letter above two columns indicates no significant difference, different letters stand for significant differences according to the conditions; B: Diagram representing percentage of apoptotic *Hydra* cells relative to numbers of epithelial cells from polyps after 7 days without food; Animals of the founder strain AEP (grey), transgenic animals expressing GFP targeted to mitochondria in ectodermal cells (mitoGFPecto, green) and animals expressing GFP tagged HyBcl2-like 4 in ectodermal cells (HyBcl-2-like 4, red) were analysed. Standard deviation from counts of four biological replicates; letters above columns indicate significant differences between apoptotic numbers in starved animals between AEP and transgenic lines-b above red column indicates that HyBcl-2-like 4 line counts differ significantly from counts in control animals (AEP and mitoGFPecto).

We then induced apoptosis in *Hydra* cells by depriving the animals of food for 7 days. Whereas regularly fed mitoGFP animals and HyBcl-2-like 4-GFP animals have approx. 1 % of apoptotic epithelial cells (Fig. 2A, 0 Wortmannin), in starved mitoGFP-control animals 5% of ectodermal epithelial cells were apoptotic. In the HyBcl-2-like 4 strain, this number was reduced to 2% (Fig. 2B) indicating that HyBcl-2-like 4 protects cells from apoptosis that is induced by starvation.

We wondered whether this protection from apoptosis in starved HyBcl-2-like 4 animals could allow budding that was normally suppressed by food deprivation. This was not the case. When we compared budding rates of HyBcl-2-like 4 - and control polyps budding rates were the same in all strains (Fig. 3).

**Figure 3:**
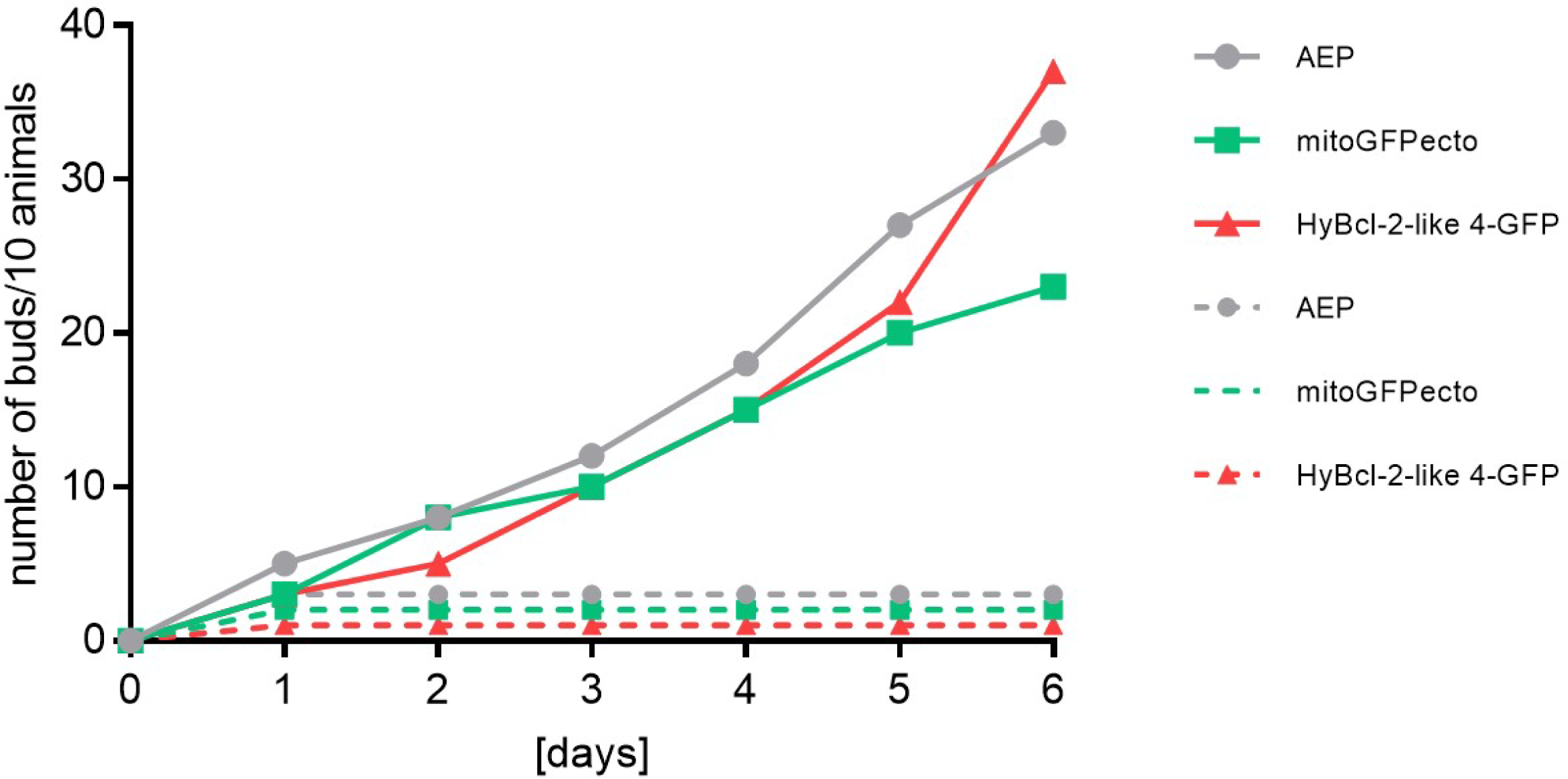
Budding rates of *Hydra* polyps in fed and non-fed animals over 6 days; animals of the founder strain AEP (grey) and transgenic animals mitoGFPecto (green) and HyBcl2-like 4 (red) were analysed. Straight lines indicate fed animals, dotted lines indicate starved animals.

Taken together, these results confirm an anti-apoptotic role for HyBcl-2-like 4 in *Hydra* as it had been suggested from our previous experiments expressing HyBcl-2-like proteins in human cells. Moreover, apoptosis was also reduced under physiological conditions of starvation, indicating a generally conserved function of anti-apoptotic Bcl-2 family members in animals from pre-bilaterians to humans. In addition, apoptosis does not seem to function as a major signal that regulates budding in response to nutrient supply in *Hydra.* However, as the transgene was only present in ectodermal cells this requires further investigation.

### The Hydra Lifeguard and Bax-Inhibitor genes

We then investigated *Hydra* genes encoding members of the TMBIMP family of cytoprotective proteins that could play a role in regulation of apoptosis independently of HyBcl-2-like proteins. We searched the *Hydra* genome and ESTs (Expressed sequence tags) for homologs and found one Bax-Inhibitor homolog (HyBI-1) and three Lifeguard homologs (HyLfg-1a, HyLfg-1b and HyLfg-4). Full-length cDNAs of these four genes were cloned and sequenced. Phylogenetic analysis of HyLfgs, HyBI-1 and their homologs from vertebrates, plants, *Caenorhabditis elegans, Drosophila melanogaster* and *Nematostella vectensis* was performed. The analysis was based on the maximum likelihood (ML) method. The tree in Fig. 4 shows two main branches separating Bax-Inhibitor (TMBIMP6, blue branch) and Lfg-4-sequences (TMBIMP4, green branch for plants; red branch for animals) from all other animal Lfg-sequences, including Lfg-5 (TMBIMB1b, lila branch), Lfg-1 (TMBIMB3, grey branches including invertebrate and vertebrate sequences), Lfg-2 (TMBIMP2, pink branch)) and Lfg-3 (TMBIMP1, brown branch). BI-1 sequences are found in animals and plants and this is also the case for Lfg-4 sequences, where the plant sequences have expanded. In animals we only find one Lfg-4 sequence in each species that we have included in the analysis. Cnidarian genes (*Hydra* and *Nematostella*) are clearly represented in the animal branches of both, BI-1 and Lfg-4 genes. In contrast, Lfg-4 is not found in the genome of *Caenorhabditis elegans* (Henke et al., 2011).

**Figure 4:**
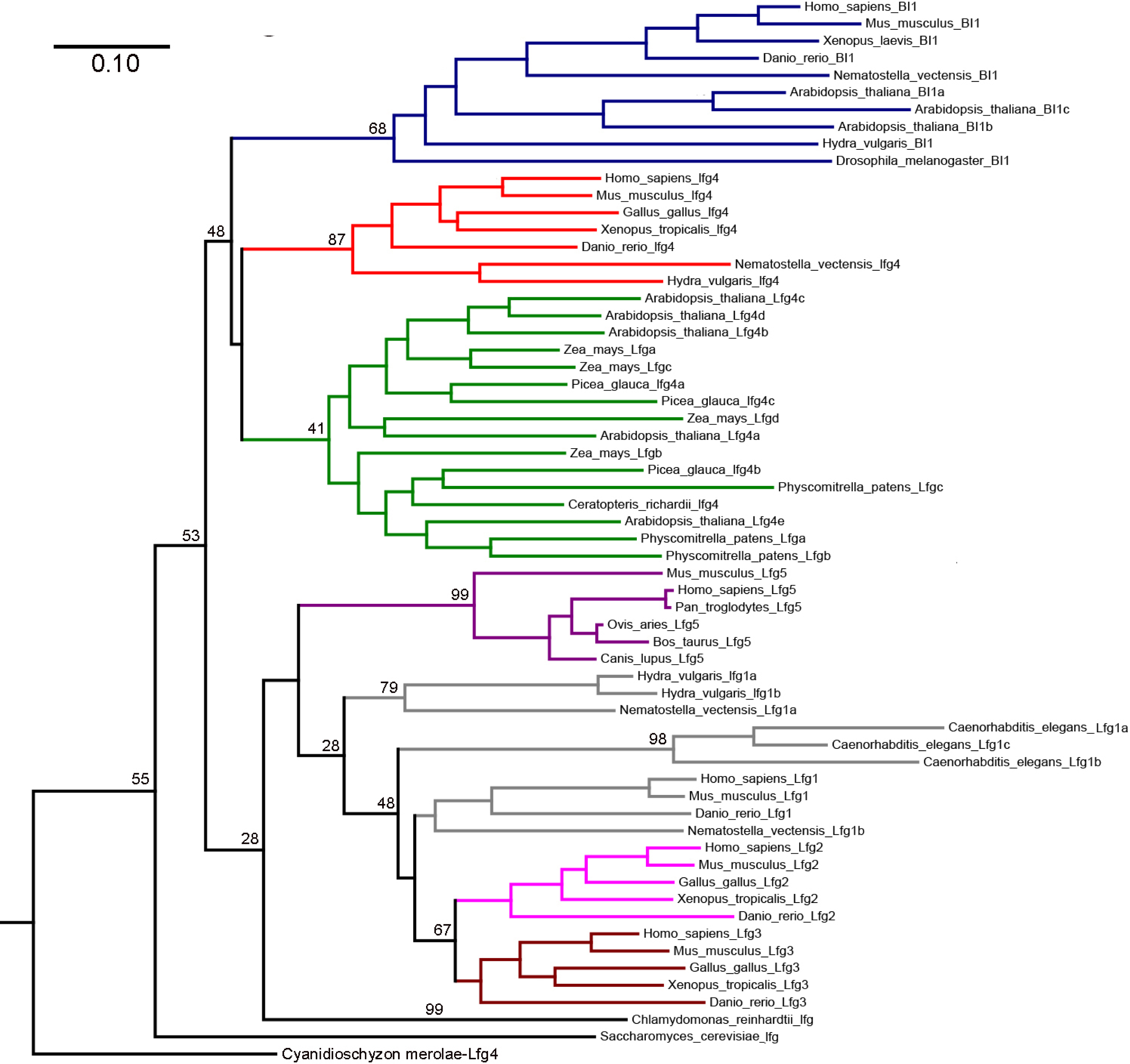
Maximum likelihood phylogenetic tree from *Muscle* alignment including TMBIMP-family members BI-1 (Blue branches), Lfg-4 from animals (red branches) and plants (green branches), Lfg-1 (grey branches), Lfg-2-(pink branches), Lfg-3 (brown branches) and Lfg-5 (purple branches) family members from indicated species including green and red algae, *Saccharomyces cerevisiae*, animals and plants. Bootstrap probabilities from 10.000 iterations are given.

The sequences for Lfg-1, -2, -3 and -5 appear as a sister branch of the Lfg-gene of the green algae *Chlamydomonas reinhardtii*. The *Hydra* sequences are clustered with *Nematostella* and *Caenorhabditis* sequences and separated from Lfg-1, -2 and -3, which are present in higher animals. Lfg-5 and Lfg-1 have been considered by other authors to be ancestral to Lfg-2 and Lfg-3 derivatives (Mariotti et al., 2014). The *Hydra* Lfg-1a and -1b do not cluster with Lfg-5, which therefore could have been lost in this lineage.

Our phylogenetic analysis and the placement of the *Hydra* sequences is in accordance with the hypothesis that the TMBIMP gene family expanded from a single ancestor that arose before the divergences of animals, plants, fungi and protozoa (Hu et al., 2009). It then split into BI-1 and Lfg-4 lineages, the latter expanded in plants. In animals a Lfg-1/5 like sequence evolved and expanded in both, invertebrates and vertebrates. It also got lost from some phyla (e.g. birds and reptiles (Hu et al., 2009; Mariotti et al., 2014)).

Closer analysis of the HyBI-1 and HyLfg sequences provides additional confirmation for the grouping of these sequences within the phylogenetic tree. Fig. 5A shows an alignment of HyBI-1 with human BI-1. The Bax-Inhibitor motif (PS01243) responsible for the name of the TMBIMP-family (see above) is labelled in bold red letters and, although clearly recognisable, it is not completely identical between the *Hydra* and human sequences. For the HyBI-1 sequence the TMHMM program predicts a six α-helix transmembrane domain structure with intracellular C- and N-terminal regions (Fig. 5A), similar to human BI-1. Moreover, the C-terminal region of BI-1 is also conserved between *Hydra* and *Homo sapiens*. It contains a semi-hydrophobic helix with four conserved aspartate residues and a very basic C-terminal region, indicated in tetramerisation of the protein and in pH-sensing (Bultynck et al., 2012; Henke et al., 2011)(Fig. 5A).

**Figure 5:**
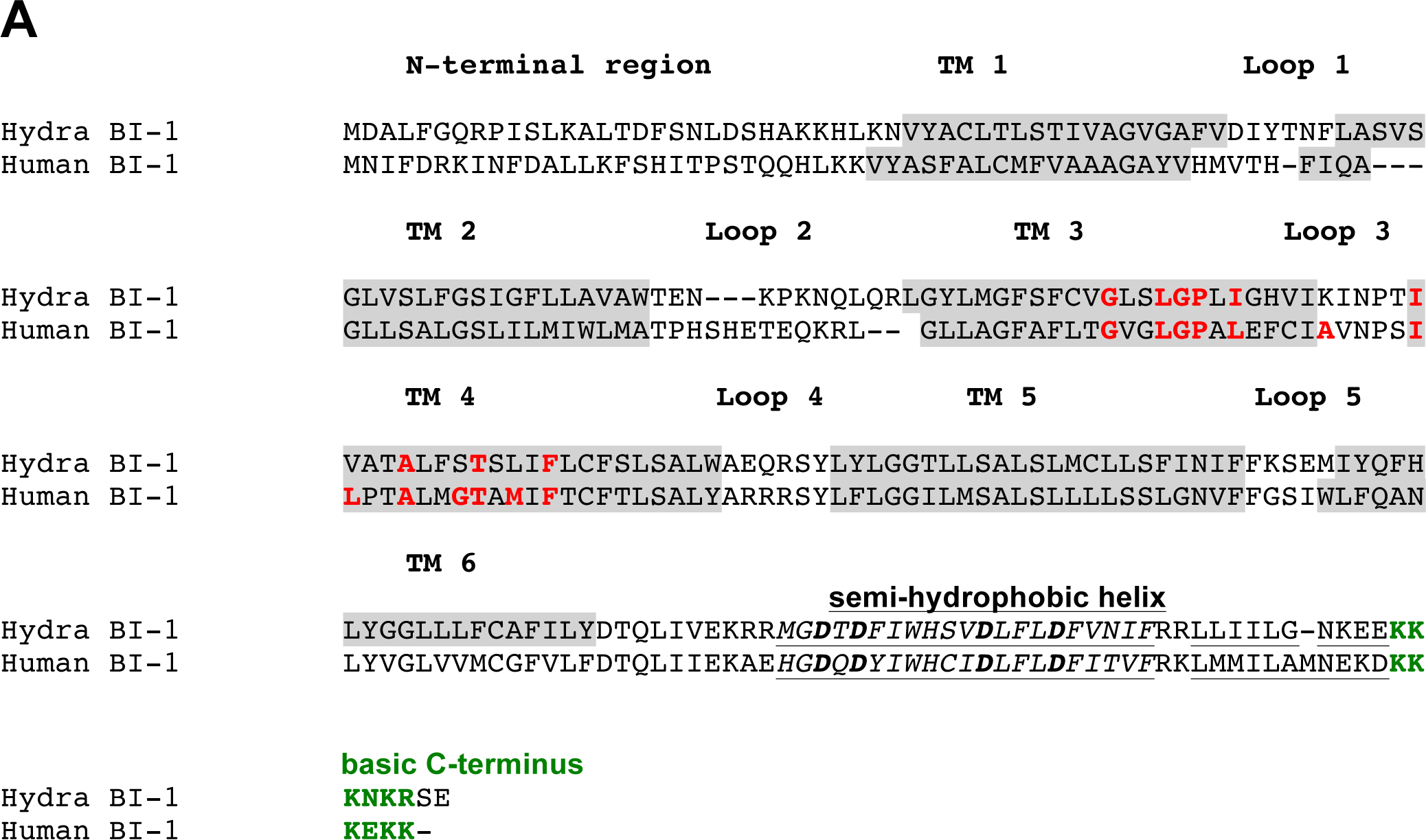
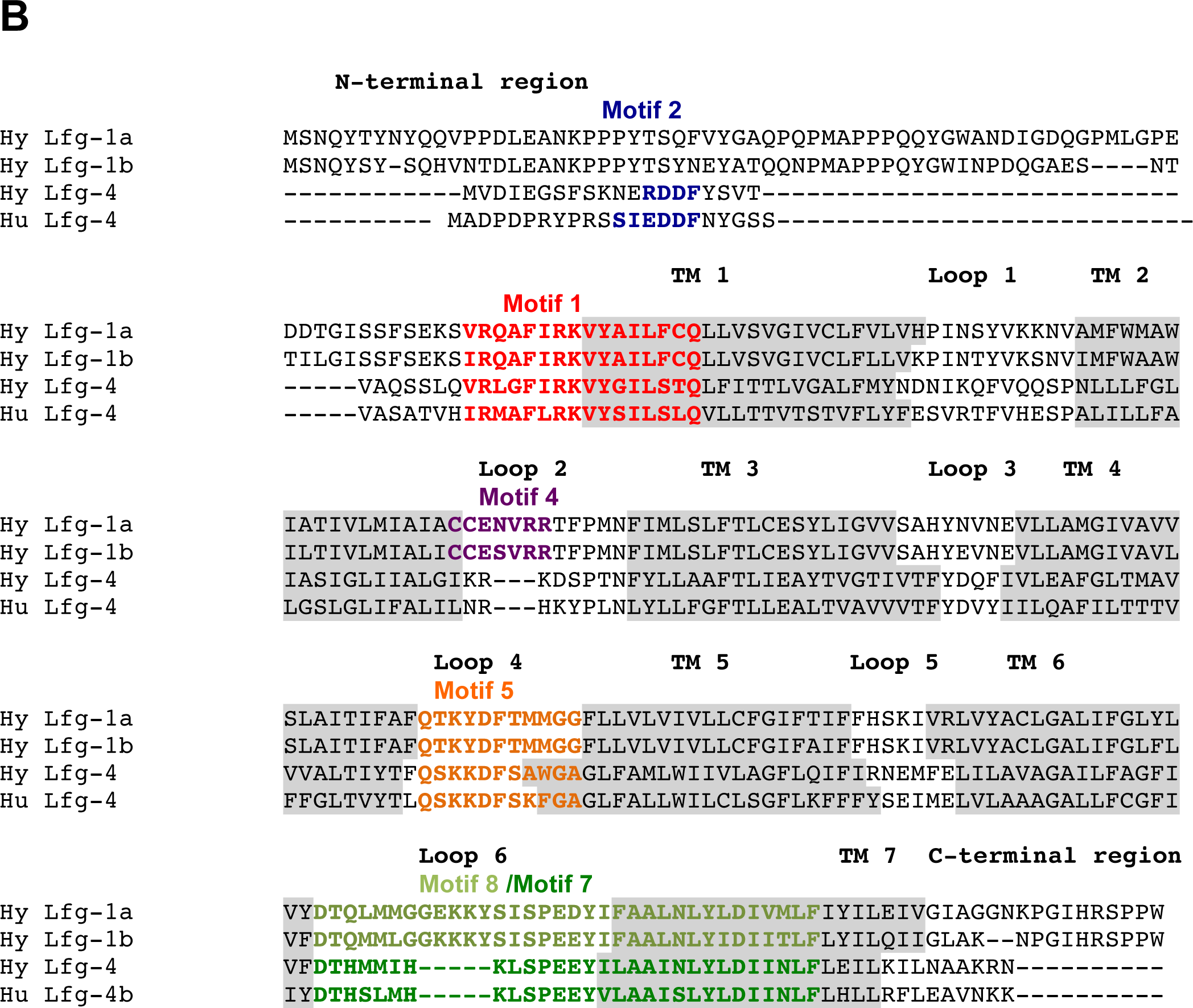
A: Alignment of HyBI-1 and human BI-1 protein sequences; Transmembrane domains from TMHMM predictions in grey boxes, transmembrane domains (TM) and loop regions (Loop) are numbered and indicated above the sequences. The predicted semi-hydrophobic helices are in *italics* with arginine residues in **bold** and the C-terminal basic domain is illustrated by green letters. The consensus amino acids of the Bax-Inhibitor motif including TMs 3 and 4 and the loop 3 sequences are labelled by red letters. B: Alignment of HyLfg-1a, HyLfg-1b, HyLfg-4 and human Lfg-4 (isoform 2). Transmembrane domains from TMHMM predictions in grey boxes, transmembrane domains (TM) and loop regions (loop) are numbered and indicated above the sequences, conserved motifs in Lfg-proteins according to (Hu et al, 2009) are indicated by blue, red, orange and green letters.

Fig. 5B shows an alignment of all three *Hydra* Lfg-proteins with human Lfg-4 (isoform b). The TMHMM program predicts a seven α-helix transmembrane domain structure with an extracellular C-terminal region for all *Hydra* proteins, similar to human Lfg-4. Ten characteristic sequence motifs diagnostic of the Lifeguard family have been described (Hu et al., 2009). Fig. 5B shows that the appropriate diagnostic motifs are conserved in HyLfgs. Motif 1 in the N-terminal and transmembrane region 1 (TM 1) is present in all Lfg-family members in animals and in plants and is also present in the three *Hydra* Lfgs. Motif 2 is only found in Lfg-4 proteins in animals and accordingly we have identified it in *Hydra* Lfg-4. Motifs 3 and 6 are specific for Lfg-2, -3 and -5 family members in animals and they are not found in any of the *Hydra* Lfgs. In contrast, motif 4, which is specific for animal Lfg-1, -2, -3 and -5 is present in HyLfg-1a and HyLfg-1b within the predicted second intracellular loop. Motif 5 in the fourth intracellular loop is conserved in all animal lifeguard members and, accordingly in all *Hydra* family members. Motifs 7 and 8 are very similar and they are distinguished with respect to their difference in animal Lfg-1, -2, -3 and -5 (motif 8) and Lfg-4 (motif 7). They are present in the *Hydra* proteins with motif 8 in Lfg-1a and-1b and motif 7 in Lfg-4. Motifs 9 and 10 belong to the plant Lfgs and have not been identified in animals and, thus they were not found in HyLfgs.

In summary, as has been indicated by previous research, the TMBIMP family is highly conserved and in animals clearly divided in BI-1, Lfg-4 and Lfg-1, 2, 3 and 5 groups, whereby *Hydra* and other earlier derived animals only have genes related to Lfg-1. The originally assigned BI-1 motif (PS01243) in *Hydra* is specific for BI-1 like family members (see also (Hu et al., 2009), Lfg-family members possess a number of highly conserved motifs that are not present in BI-1 and show a specific signature in different Lfg-proteins between animals and plants and between Lfg-4 and animal specific Lfg-1, -2, -3 and -5 proteins. As we show here, specific sequence motifs for animal Lfg-1 related proteins are present in *Hydra* and therefore have evolved before the divergence of bilaterians.

### Subcellular localisation of HyBI-1 and HyLfg proteins

In order to get an idea about a possible conservation of function of TMBIMP family members we next sought to investigate the subcellular localisation of these proteins in *Hydra*. Therefore, we transfected *Hydra* polyps with plasmids encoding GFP-HyBI-1, GFP-HyLfg-4, GFP-HyLfg-1a and GFP-HyLfg-1b proteins under the control of the *Hydra-*actin promoter in the vector HotG (Böttger et al., 2002) using a particle gun.

GFP-HyBI-1 was found in a net-like distribution all over the cytoplasm of transfected epithelial cells reminding of an ER-distribution (Fig. 6A). Co-staining with the mitochondrial α-ATPase antibody did not reveal any co-localisation (not shown). We could not find any specific marker to label the ER in *Hydra* cells. Therefore, we resorted to human cells and expressed the HyBI-1-protein fused with GFP in HEK-cells (human embryonal kidney) using the plasmid pEGFP:HyBI-1. Co-staining of transfected cells with anti-Calnexin antibody clearly showed ER-localisation of HyBI-1 in human cells (Fig. 6B).

**Figure 6:**
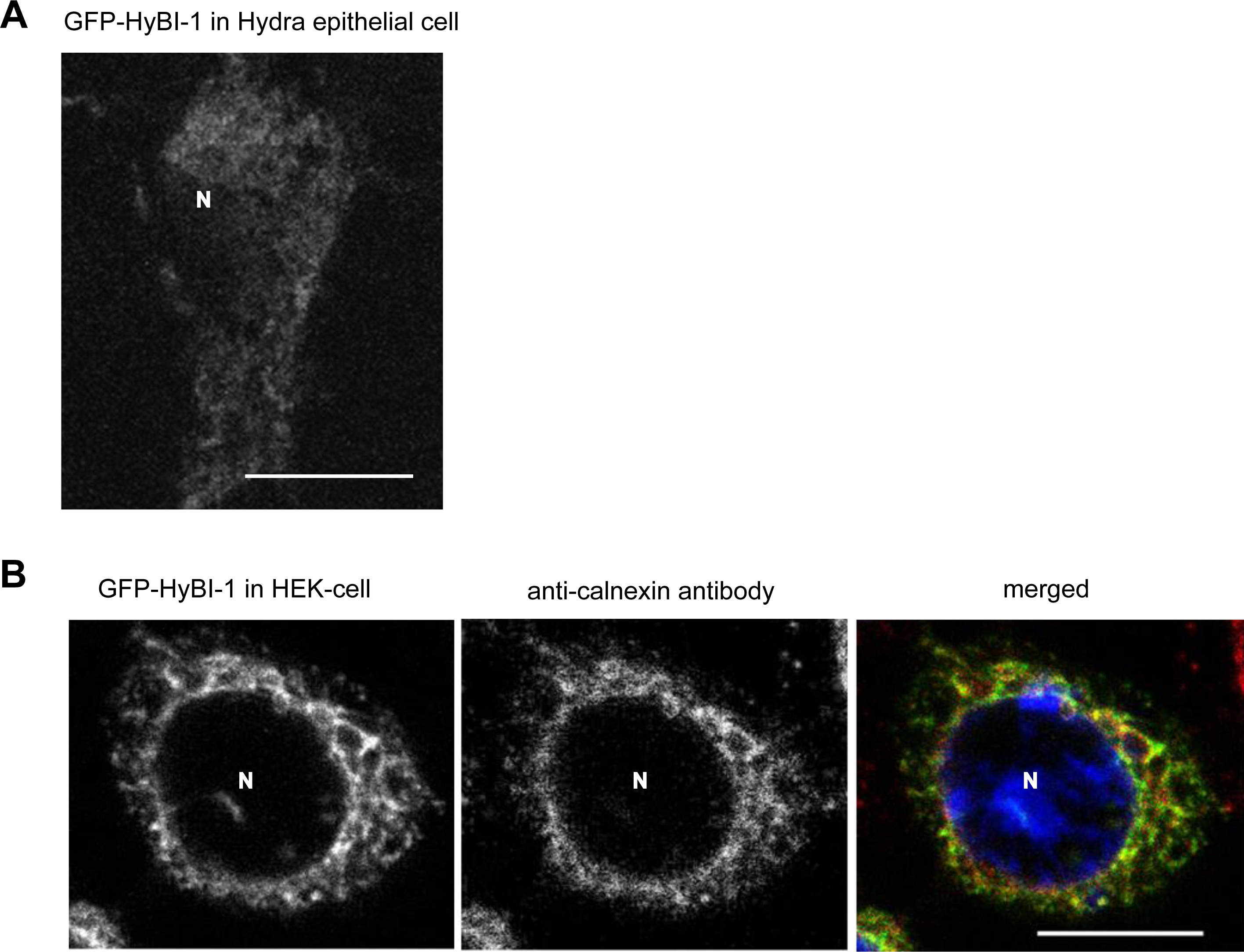
A: *Hydra* ectodermal epithelia cell expressing GFP-tagged HyBI-1 after transfection of polyps with particle gun; single section from confocal laser microscope showing GFP-signal; B: Human HEK293-cell expressing GFP tagged HyBI-1 from the plasmid pcDNA3, counterstained with anti-calnexin antibody indicating ER-localisation of HyBI-1 in human cells; single sections from laser confocal microscope and merged images are shown, DAPI-image blue and only in merged image, N indicates cell nucleus. Scale bars 10 µm

GFP-HyLfg-1a and GFP-HyLfg-1b were found localised at the plasma membrane and in vesicle-like structures in the cytoplasm of *Hydra* epithelial cells (Fig. 7A, B). GFP-HyLfg-4 was found localised in globular structures of *Hydra* epithelial cells but not on the plasma membrane. The Golgi- and plasma membrane marker BODYPY-TR ceramide (InVitrogen) strongly stained plasma membranes of living *Hydra* cells. Golgi vesicle staining appeared weaker. However, comparison of GFP-signals of Lfg-1a, Lfg-1b and Lfg-4 with the Golgi-and plasma membrane marker BODIPY-TR ceramide clearly revealed localisation of all three GFP-fusion proteins with these Golgi-structures and in addition co-localisation of HyLfg-1a and 1b with plasma membranes (Fig. 7A, B, C). To confirm the Golgi-distribution of BODYPY-TR in *Hydra* we used the Golgi marker on human HEK-cells and additionally transfected them with the plasmid pEGFP:HyLfg-4. Very similar to the distribution of GFP-HyLfg-4 in *Hydra*, the protein was found in a vesicle complex near the nucleus. This structure was also stained with the Golgi-marker (Fig. 7D). From this we conclude that BODYPY-TR is a suitable Golgi-marker in *Hydra* and that Hy-Lfg-4 is a Golgi-protein.

**Figure 7:**
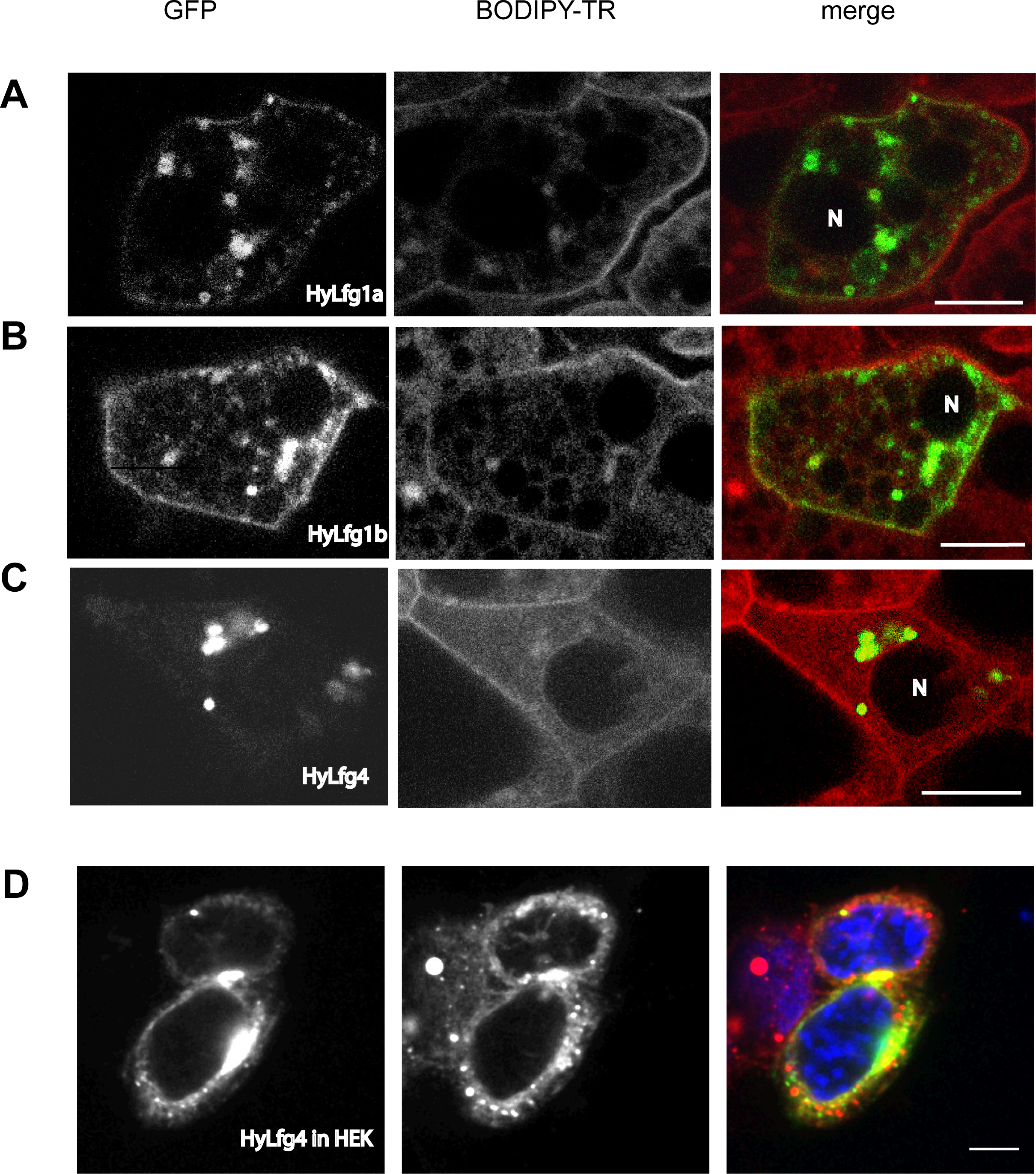
A-C: Confocal microscopic single optical sections of *Hydra* ectodermal epithelial cells of live animals, A: expressing GFP-tagged HyLfg-1a; B: expressing GFP-tagged HyLfg-1b; C: expressing GFP-tagged HyLfg-4, all are counterstained with BODIPY-TR ceramide (InVitrogen, middle panels). Merged images in right hand panels, N indicates nucleus; D: Human HEK-cell expressing GFP-tagged HyLfg-4, counterstained with BODIPY-TR ceramide, Scale bars A-C: 10 µm; D: 5 µm

### HyBI-1 protects human cells from apoptosis

The ER-localisation of HyBI-1 as well as the high sequence conservation extending into the C-terminal domain prompted us to ask whether the HyBI-1 protein was involved in mechanisms of programmed cell death. Therefore, we overexpressed GFP-HyBI-1 protein in HEK-cells and counted the percentage of apoptotic cells in relation to transfected cells that showed a GFP-signal. HyBI-1 did not induce apoptosis above the level found in control cells. However, it was able to protect cells from camptothecin induced apoptosis. The topoisomerase inhibitor camptothecin induced apoptosis in 70 % of control cells (overexpressing GFP) within 24 hours of application (Fig. 8). This effect was greatly diminished in cells overexpressing GFP-HyBI-1 indicating that HyBI-1 strongly protected HEK-cells from apoptosis.

**Figure 8:**
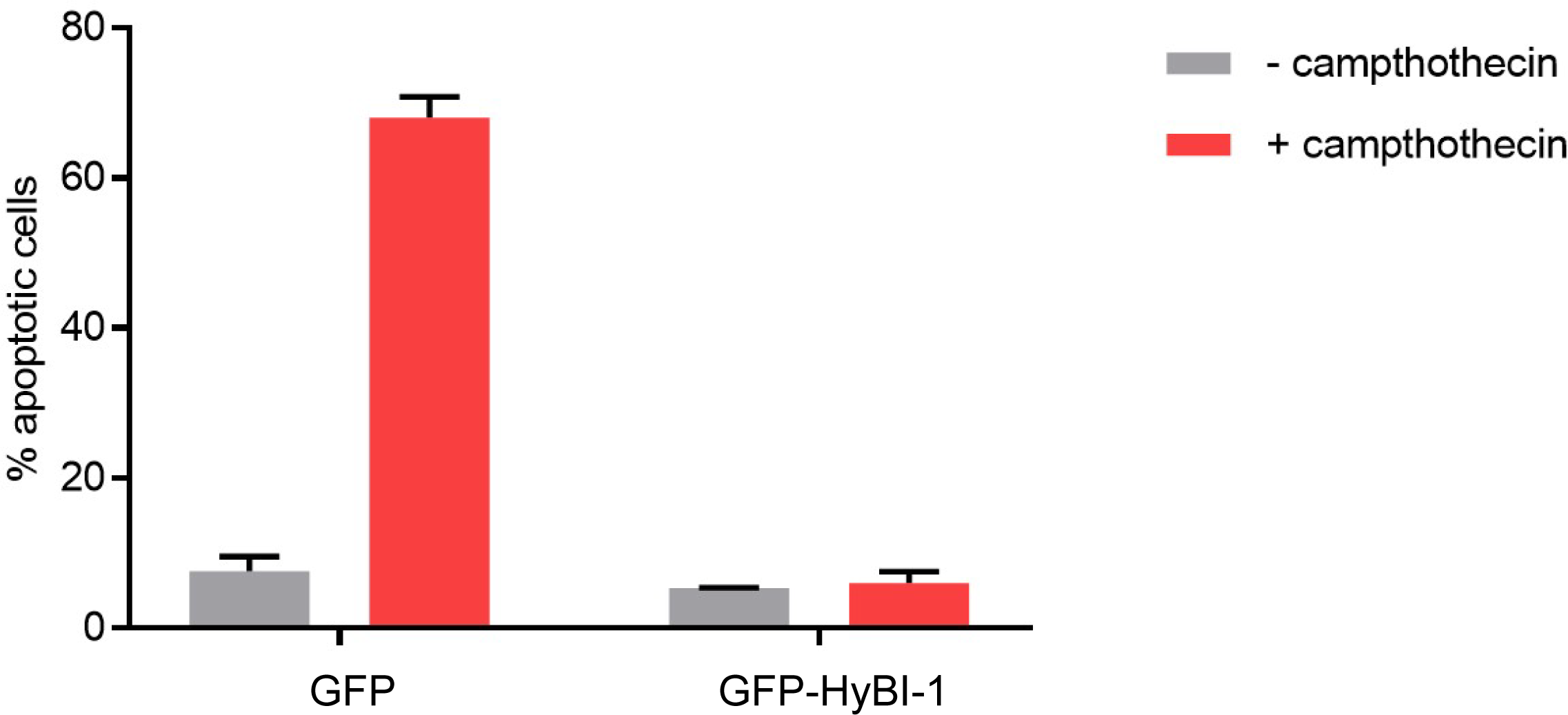
Diagram displaying result of apoptosis assay in HEK-cells; GFP or GFP-tagged HyBI-1 were expressed in HEK-cells. GFP-positive apoptotic cells as percentage of all GFP-positive cells were counted in untreated cells (-camptothecin) and in cells treated with camptothecin (+camptothecin).

### HyBI-1 and HyLfg expression

We next wanted to investigate to which levels *Hydra* TMBIMPs were expressed and whether their expression responded to apoptotic stimuli. At first we estimated the expression levels of TMBIMP-genes by RT-qPCR (Fig. 9). This clearly indicated that HyBI-1 is expressed at the highest level, followed by HyLfg-4 with an 8 times lower expression level. Lfg-1a and Lfg1-b as well as Bcl-2-like 4 and, for comparison HyBcl-2-like 5, are only weakly expressed. We tested several apoptosis inducing conditions, including benzamidine, camptothecin, UV light, Wortmannin and starvation for their effect on the transcriptional level of these genes. None of these stimuli changed gene expression significantly (not shown).

**Figure 9:**
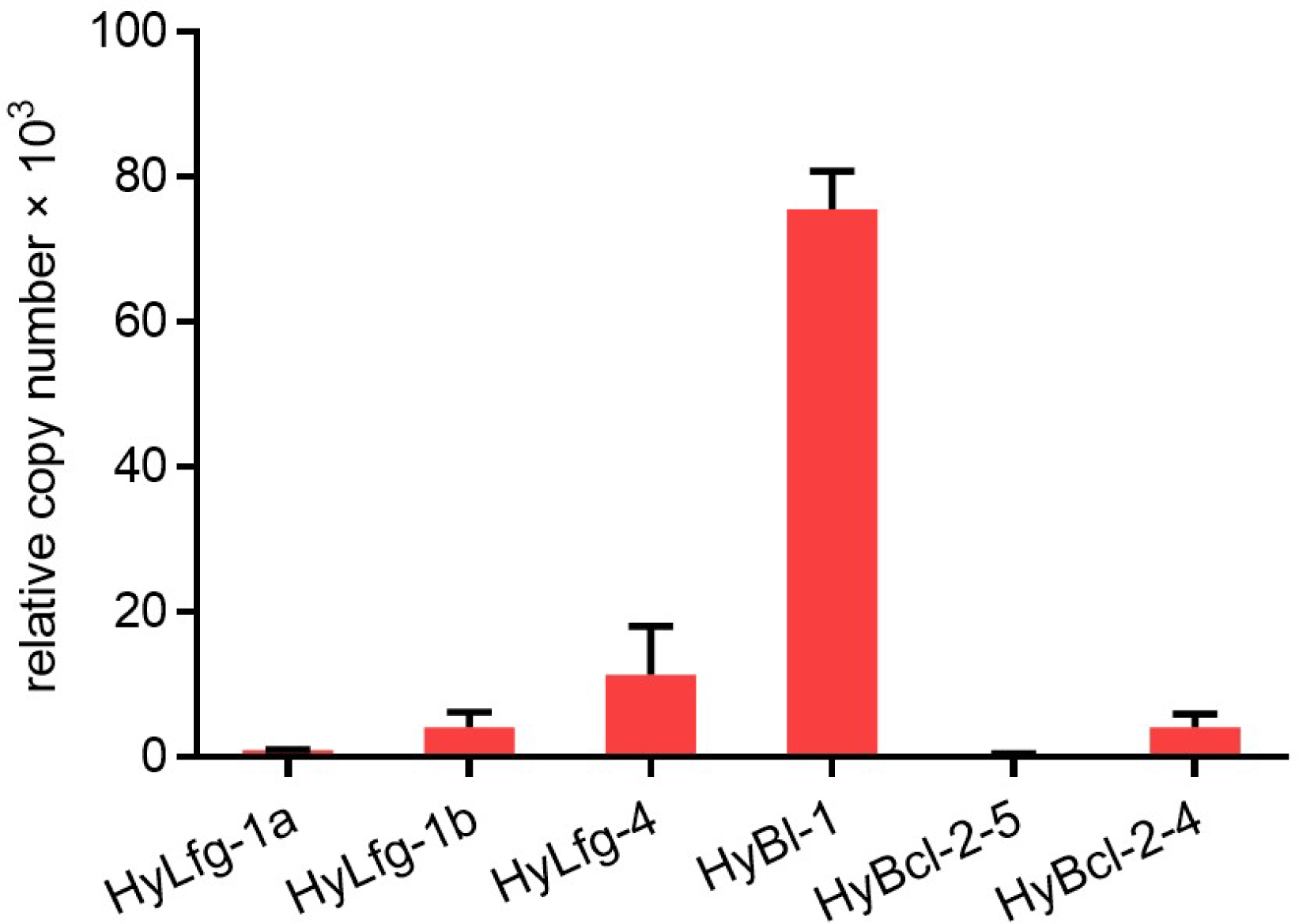
Diagram displaying relative expression levels of Bcl-2 and TMBIMP-family members in *Hydra* as determined by RT-qPCR. Relative copy number is an arbitrary unit.

## Discussion

Programmed cell death is an important developmental mechanism and it probably occurs in all organisms. The molecular mechanisms that regulate and execute this process differ between animals, fungi and plants. However, some protein families involved in regulating programmed cell death are conserved through the whole tree of life. This clearly includes the TMBIMP family of membrane proteins. On the other hand, pro- and anti-apoptotic members of the Bcl-2 family of proteins that are largely associated with mitochondrial membranes, are only conserved in animals. They have neither been found in plants, nor fungi. In cnidarians, one of the very first derived metazoan phyla, they already form a large family. This is all the more striking when we consider that cell death in the nematode *Caenorhabditis elegans* only requires one *bcl-2* related gene, *ced-9*. In *Drosophila* two *bcl-2* genes exist, however they do not appear to be essential for apoptosis. It was therefore very important to investigate whether the elaborated Bcl-2 family in *Hydra* performs similar functions as in mammals. Our previous work had indicated that some members of this protein family can interact with each other via their BH3-domains and that they exhibit either pro- or anti-apoptotic functions in human cells. Here we now complement these data with experimental evidence for one family member, HyBcl-2-like 4, to act as an inhibitor of apoptosis in *Hydra* epithelial cells. This was evident when we induced apoptosis with the PI(3)kinase inhibitor Wortmannin in transgenic animals that overexpressed GFP-Bcl-2-like 4. Wortmannin has been shown previously to induce massive apoptotic cell death in cell types of the interstitial cell lineage and some cell death in epithelial cells in *Hydra (David et al., 2005).* From these observations we had concluded that survival of *Hydra* cells is mediated by the PI(3)-kinase-PKB-pathway and thus was most likely dependent on extracellular growth and/or survival factors. So far, these factors are not known. We had also hypothesised that survival factors could regulate the cellular response of *Hydra* to limited food supply. When the animals are starved for one week, they maintain their normal proliferation rates, although cell numbers do not increase and bud formation ceases. Thus, “excess” cells are produced which die by apoptosis (Bosch and David, 1984). Whereas regularly fed animals have an apoptotic index of ca. 1 % (see Fig. 2A, 0 Wortmannin) this increases in non-fed animals up to 5 % (Fig. 2B). In HyBcl-2-like 4 overexpressing animals this increase of apoptotic cells was strongly diminished (Fig. 2B) suggesting that HyBcl-2-like 4 also blocks starvation dependent apoptosis.

The observed decrease in apoptosis in HyBcl-2-like 4 animals did not lead to higher numbers of buds during the starving period. It is possible that this was because HyBcl-2-like 2 was only overexpressed in ectodermal cells and not in endodermal or interstitial cells. On the other hand, the regulatory circuit between cell death, cell numbers, animal size and budding (reproduction) should be investigated further on the level of signalling. It is known that extensively fed *Hydra* polyps can grow much larger than normally fed ones and then both, budding rate and animal size increase. It remains speculative at the moment whether HyBcl-2-like 4 and possibly other members of the Hydra Bcl-2 family are involved in this regulation. What we can conclude from our data is that HyBcl2-like 4 as a mitochondrial protein in Hydra fulfils a similar function in protecting cells from apoptosis as it does in mammals.

We then investigated the TMBIMP protein family in *Hydra*. Some members of this family had been implicated in cytoprotection in plants and in animals (Hu et al., 2009; Reimers et al., 2008; Somia et al., 1999). We cloned HyBI-1 and three HyLfg-genes from *Hydra* cDNA and performed a phylogenetic analysis. This revealed three groups comprising Lfg-1 derived proteins, Lfg-4 related proteins and Bax-Inhibitor related proteins. The *Hydra* sequences fit very clearly into these groups.

Lfg-1 is conserved from plants to humans. However, in mammals it splits into four different groups, Lfg-1, 2, 3 and 5. Invertebrate Lfg-1 proteins in *Hydra* and *Caenorhabditis elegans* also have two Lfg-1 homologs each. In a recent study the subcellular localisation of all human TMBIMPs was analysed. It was shown that human BI-1 and TMBIMP 4 and 5 (corresponding to Lfg-4 and 5) were localised in the ER, whereby TMBIMP 5 could also be sorted to mitochondria. In contrast, TMBIMPs 1-3, including Lfg-1 were found associated with the Golgi (Lisak et al., 2015). Localisation studies by other authors had indicated similar subcellular distributions of TMBIMP family proteins, except for Lfg-4, which was found associated with Golgi membranes in human cells and it was therefore called Golgi-associated anti apoptotic protein (GAAP) reviewed in (Carrara et al., 2017). In comparison with *Hydra* and consistent with the studies in human cells, HyBI-1 was localised in the ER and HyLfg-1a and -1b were associated with Golgi-vesicles. HyLfg-4 is found in Golgi vesicles in *Hydra* (Lisak et al., 2015).

To gain some insight into a possible physiological function of BI-1 and Lfg-proteins in *Hydra*, we carried out expression studies. We found that Lfg-1a and -1b were not expressed at high levels in *Hydra*. This was also true for the pro-apoptotic *Hydra* Bak-1. On the other hand, HyBI-1 was expressed at high levels and HyLfg-4 had a middle strong expression level in non-sexual *Hydra*.

HyBI-1 distribution on the cellular level by excluding the nuclei and resembling a reticular structure in the cytoplasm looks ER-like. We could not provide or find any agent suitable to serve as a marker for ER in Hydra. However, taking into account the very clear co-localisation of HyBI-1 with calreticulin after overexpression of the protein in human cells and the KDEL-signal present in the *Hydra* protein suggests that HyBI-1 is also localised in the ER in *Hydra* where we now even would recommend to use HyBI-1-GFP as a *Hydra-*marker for this cellular structure.

Taken together, our data indicate that both, the Bcl-2 family proteins and the TMBIMP-family proteins are involved in apoptosis that regulates tissue homeostasis and development in *Hydra*. They do not seem to be regulated significantly on the transcriptional level by external apoptotic stimuli, like DNA damage.

Many hypotheses about the molecular evolution of apoptosis focus on the involvement of mitochondria in this process. This is often connected with the endosymbiosis theory for the evolution of eukaryotic cells, whereby a symbiosis arose between a α-proteobacterium and a member of the archea (Ku et al., 2015). This could then have induced an evolutionary “battle” where the proteobacterium may have employed membrane pore forming molecules (such as Bcl-2 family members are) and injected toxins into the cytoplasm of the archaea, e.g. holocytochrome c. However, Bcl-2 like proteins are only found in animals and thus a large gap is evident between the evolution of the first eukaryotic cells and the evolution of multicellular animals that perform apoptotic cell death (recently reviewed by (Green and Fitzgerald, 2016). TMBIMP-family proteins could fulfil similar functions that are allocated to Bcl-2 family proteins in animals. Moreover, they are possibly responsible for such regulations in cellular membrane compartments other than mitochondria.

**Here we have shown that anti-apoptotic Bcl-2 family proteins and proteins of the TMBIMP-family represent mitochondrial ER-, Golgi and plasma membrane proteins that fulfil cytoprotective functions in *Hydra*. These are closely intertwined with mechanisms that regulate apoptotic cell death in animals. The Bcl-2 family arose right at the beginning of metazoan evolution and performs functions in regulating cellular homeostasis in response to nutrient supply. One of the TMBIMP family members in *Hydra* is a potential apoptosis inhibitor. TMBIMP-family proteins are conserved in plants and animals and HyBI-1 localises to the ER and suppresses programmed cell death in plants (Bolduc and Brisson, 2002). Therefore, by trying to understand the evolution of molecular mechanisms of programmed cell death the focus on mitochondria could be extended and other membrane bound cellular compartments should be taken into consideration.**

## Acknowledgements

This work was funded by DFG-grant awarded to A.B BO1748-7-1.

